# Mechanisms of type I interferon production by chicken TLR21

**DOI:** 10.1101/2023.09.20.558228

**Authors:** Rodrigo Guabiraba, Damaris Ribeiro Rodrigues, Paul T. Manna, Mélanie Chollot, Vincent Saint-Martin, Sascha Trapp, Marisa Oliveira, Clare E Bryant, Brian J Ferguson

## Abstract

The innate immune response relies on the ability of host cells to rapidly detect and respond to microbial nucleic acids. Toll-like receptors (TLRs), a class of pattern recognition receptors (PRRs), play a fundamental role in distinguishing self from non-self at the molecular level. In this study, we focused on TLR21, an avian TLR that recognizes DNA motifs commonly found in bacterial genomic DNA, specifically unmethylated CpG motifs. TLR21 is believed to act as a functional homologue to mammalian TLR9. By analysing TLR21 signalling in chickens, we sought to elucidate avian TLR21 activation outputs in parallel to that of other nucleic acid species. Our analyses revealed that chicken TLR21 (chTLR21) triggers the activation of NF-κB and induces a potent type-I interferon response in chicken macrophages, similar to the signalling cascades observed in mammalian TLR9 activation. Notably, the transcription of interferon beta (*IFNB*) by chTLR21 was found to be dependent on both NF-κB and IRF7 signalling, but independent of the TBK1 kinase, a distinctive feature of mammalian TLR9 signalling. These findings highlight the conservation of critical signalling components and downstream responses between avian TLR21 and mammalian TLR9, despite their divergent evolutionary origins. These insights into the evolutionarily conserved mechanisms of nucleic acid sensing contribute to the broader understanding of host-pathogen interactions across species.

## 1. Introduction

The ability of host cells to rapidly sense and respond to microbial nucleic acids is an essential part of the innate immune response to infection (Bryant et al., 2015; Pandey et al., 2014). Endosomal Toll-like receptors (TLRs), a family of pattern recognition receptors (PRRs), bind directly to multiple specific DNA and RNA species (Majer et al., 2017). PRRs evolved as one mechanism by which organisms can distinguish self from non-self at the molecular level. This is achieved in the context of nucleic acid sensing by recognition of specific nucleic acid structures or motifs that are not normally found in host cells and tissues under homeostatic conditions. In the cytoplasm, PRRs such as cGAS and DNA-PK pathway sense and respond to viral dsDNA in a DNA length-dependent but DNA-sequence-independent manner (Bryant et al., 2015; Oliveira et al., 2020). On the other hand, nucleic acids taken up into endosomes are sensed by TLRs. In this context, DNA that contains unmethylated CpG motifs is rare in vertebrates due to high levels of chromatin methylation, but common in bacterial genomic DNA (Krieg et al., 1995). CpG DNA therefore acts as a pathogen-associated molecular pattern (PAMP) and several TLRs have evolved to directly bind DNAs that contain this motif. In vertebrates there are two TLRs known to directly bind CpG-containing DNAs; TLR9 (Hemmi et al., 2000) and TLR21 (Chuang et al., 2020; Keestra et al., 2010). These TLRs have different species distributions and a different evolutionary root (Temperley et al., 2008), despite their ability to bind similar, if non-identical, ligands (Yeh et al., 2013). One striking difference between these TLRs is the absence of TLR21 in mammals, where TLR9 is the predominant CpG sensor, and the absence of TLR9 in birds, where TLR21 is the only recorded CpG sensor. Chicken TLR21 is the best characterised avian TLR21. chTLR21 responds to a range of CpG DNA oligonucleotides (ODN), senses bacteria including *Campylobacter* spp. and triggers downstream signalling (Brownlie et al., 2009; de Zoete et al., 2010; Keestra et al., 2010). Given the extensive use of CpG DNAs as immunomodulatory agents, understanding the signalling mechanisms and outputs of TLR21 stimulation has relevance not only in the context of antimicrobial responses but also in vaccine and immunoregulatory pharmaceutical design.

The signalling outputs of TLRs include the transcription of inflammatory mediators and interferons, and the initiation of programmed cell death (Bryant et al., 2015). From inside the endosomal compartment, nucleic-acid sensing TLRs signal through the membrane into the cytoplasm to start a signalling cascade. Endosomal nucleic acid sensing TLRs can activate NF-κB signalling via recruitment of MyD88 and activation of IRAK kinases. Some endosomal TLRs, such as TLR3, can also recruit the adaptor protein TRIF and drive type-I interferon production and regulated cell death via recruitment of the kinases TBK-1/IKKε or RIPK1/3 respectively (Bryant et al., 2015). In mammals, TBK-1 activity downstream of TLR3 results in the activation of the transcription factor IRF3. On the other hand, TLR9 ligation with CpG DNA in mammals can result in an alternative pathway that results in IRF7 activation, independent of TBK1 and IRF3 (Honda et al., 2005a). Although NF-κB activation is one output of TLR21 signalling (Keestra et al., 2010), it is not clear whether these independent pathways function downstream of TLR21 in birds. One reason for this is the loss of IRF3 in most bird species, including chickens (Santhakumar et al., 2017). In place of IRF3, there is an orthologue of mammalian IRF7, present in chickens, that functions downstream of several PRRs (Cheng et al., 2019; Santhakumar et al., 2017), although it is not clear whether chIRF7 can substitute for all mammalian IRF3 functions and this has not been studied in the context of TLR signalling in birds. There is a tight regulation of vertebrate IRF7 expression, and its activity is imperative in dictating appropriate type I IFN production for normal IFN-mediated physiological functions. IFN-I is one of the most effective antiviral innate immune mediators. Secretion and subsequent ISG transcription induced by autocrine and paracrine IFN receptor signalling sets an antiviral state in infected and bystander cells. As an example, chicken IFNβ was shown to be an autocrine/paracrine pro-inflammatory mediator in chicken macrophages, with direct effects in macrophage effector functions (Garrido et al., 2018; Oliveira et al., 2020).

We therefore set out to analyse the activity and function of TLR21 in chickens and to determine which of the known TLR-driven signalling mechanisms function downstream of this receptor. Using a full *in vitro* heterologous system comprising WT, KO, and reporter chicken cells, we have determined that chTLR21, as well as activating NF-κB, drives a potent type-I interferon response in chicken macrophages. The ability of chTLR21 to induce the transcription of interferon beta is dependent on both NF-κB and IRF7 signalling, but independent of TBK1. In addition, we show that this receptor does not directly activate cell death signalling pathways. Hence the signalling downstream of chTLR21 is similar to mammalian TLR9 signalling in its capacity to activate NF-κB and IRF7 and to produce a potent inflammatory and type-I interferon response in myeloid cells.

## 2. Materials and Methods

### 2.1 Sequence retrieval and phylogenetic analysis

Putative avian homologues of genes of interest were retrieved by BLASTp from a panel of 46 broadly representative genomes sampling avian diversity. Bird genomes searched are listed in **Supplementary Table 1**. Further searches were carried out in representative amphibian (*Xenopus tropicalis*), fish (*Danio rerio*, *Ictalurus punctatus*, *Cyprinus carpio*, *Esox Lucius*), reptile (*Gopherus agassizii, Psuedonaja textilis, Podarcis muralis, Chelonoidis abingdonii, Terrapene triunguis*), and mammalian (*Mus musculus, Rattus norvegicus*) genomes. For IRF searches, human IRF3 and IRF7 were used as query sequences. For TLR searches, chicken, xenopus, zebrafish, and mouse sequences were used as queries as appropriate. Putative homologues were initially assessed by reciprocal best hit BLASTp analysis against the genome of the respective query sequence. For phylogenetic analysis, sequences were aligned with MAFFT (version 7.110) using L-INS-I preset (Katoh and Standley, 2013). Regions containing phylogenetically informative sites were identified within aligned sequences using TrimAI (version 1.3) (Capella-Gutiérrez et al., 2009). Maximum likelihood phylogenetic reconstruction was carried out using IQ-TREE (Nguyen et al., 2014). Substitution models were identified with ModelFinder, implemented within IQ-TREE. Branch support was assessed by the SH-aLRT test and the ultrafast bootstrap approximation (Anisimova et al., 2011).

### 2.2 Cells and reagents

Calf Thymus (CT) DNA (Sigma), Herring Testes (HT) DNA (Sigma), 2’3’-cGAMP (Invivogen) and chicken interferon alpha (Yeast-derived Recombinant Protein, Kingfisher Biotech, Inc) were diluted in nuclease-free water (Ambion, ThermoFisher). The Class B CpG ODN2006 (ODN 7909, PF_3512676, sequence: 5’-tcgtcgttttgtcgttttgtcgtt-3’, Invivogen) was diluted in sterile endotoxin-free water. TLR3 agonist polyinosinic-polycytidylic acid - Poly(I:C) and TLR4 agonist LPS (Lipopolysaccharide from *E. coli* O55:B5) (Invivogen) were diluted in sterile endotoxin-free physiological water (NaCl 0.9%). BAY-11-7082, a selective and irreversible inhibitor of cytokine-induced IκBα phosphorylation, and BX795, a potent inhibitor of the noncanonical IkB kinases TANK-binding kinase-1 (TBK1) and IκB kinase-ε (IKKε) (Invivogen) were diluted in DMSO, following the manufacturer’s protocols.

HD11 cells were incubated at 37°C, 5% CO2. They were grown in RPMI (Sigma-Aldrich, Germany) complemented with 2.5% volume per volume (v/v) heat-inactivated foetal bovine serum (FBS; Sera Laboratories International Ltd), 2.5% volume per volume (v/v) chicken serum (New Zealand origin, Gibco, Thermo Fisher Scientific), 10% Tryptose Phosphate Broth solution (Gibco, Thermo Fisher Scientific), 2 mM L-glutamine (Gibco, Thermo Fisher Scientific), 50 µg/ml of penicillin/streptomycin (P/S; Gibco, Thermo Fisher Scientific).

### 2.3 Stimulation Assays

HD11 (WT, chTLR21 and chIRF7 knockouts, or reporter cells) were seeded in 6-well plates at a density of 7×10 ^5^ cells/well. In the following day, the cells were stimulated with ODN2006 (1-10 µg/ml), Poly(I:C) (5 µg/ml) or LPS (5 µg/ml), or transfected using TransIT-LT1 (Mirus Bio, USA) with HT-DNA (5 µg/ml), CT-DNA (5 µg/ml) or cGAMP (5 µg/ml). Cell samples and lysates were then harvested at different time-points post stimulation or transfection for further downstream analyses. In the NF-kB- or TBK1-inhibition assays, BAY-11-7082 (5 uM) or BX795 (3 uM) were added 1h prior to stimulation or transfection. Finally, in certain experiments, cells were pre-treated for 16h with recombinant chicken interferon alpha (50 ng/ml) prior to stimulation or transfection.

### 2.4 Reporter assays

Activation of NF-κB by different ligands was assessed using HD11-NF-κB luciferase reporter cells containing a basal promoter element (TATA box) and tandem repeats of an NF-κB consensus sequence fused to a luciferase reporter gene (Garrido et al., 2018). Cells expressing the reporter fusion are selected under puromycin selection and routinely cultured in DMEM F-12 (1:1) medium (Gibco, UK), supplemented with 10% heat-inactivated FCS, 15 mM HEPES, 2 mM L-glutamine, 100 U/ml penicillin, 100 μg/ml streptomycin and 5 μg/ml puromycin (Sigma-Aldrich, UK), and incubated as described above.

### 2.5 CRISPR gene editing

Single guide (sg)RNA sequences were designed targeting the start of the open reading frames for TLR21 (*TLR21*) and IRF7 (*IRF7*). Genome editing of HD11 was performed using ribonucleoprotein (RNP) delivery. tracrRNA was mixed with the target specific sgRNAs and, to form the RNP complex, the tracrRNA/sgRNA mix was incubated with Cas9 protein (IDT, Leuven, Belgium) and electroporation enhancer at 21°C. To generate knockout cells, 1×10e6 cells per guide were electroporated with the corresponding RNP complex using Lonza Electroporation Kit V (Lonza). After 48 h, the cells were expanded for future experiments and their DNA were extracted using the PureLink Genomic DNA Kit (Thermo Scientific, Waltham, MA, USA). The knockout efficiency was evaluated by genotyping the polyclonal cell populations using MiSeq (Illumina). Successfully edited populations were diluted to a concentration of 0.5 cell/well and seeded in 96-well plates. Individual clones were sequenced by MiSeq and the confirmed knockout clones were expanded for experiments.

### 2.6 Gene expression

Total RNA from cell lysates was extracted using the NucleoSpin RNA II kit (Macherey-Nagel, Germany) according to the manufacturer’s instructions, including an rDNAse step for the elimination of contaminant DNA. RNA quality and concentration were determined using a NanoDrop (Thermo Scientific, USA). Total RNA (0.5 μg per reaction) was reverse transcribed using the iScript cDNA synthesis kit (Bio-Rad, USA). Quantitative real-time PCR (qPCR) was performed on a CFX96 machine (Bio-Rad, USA), the reaction mixture consisting of iQ SYBR Green Supermix (Bio-Rad, USA), cDNA, primers (250 nM, Eurogentec, Belgium) and nuclease-free water (Sigma-Aldrich, UK) in a total reaction volume of 20 μL. qPCR data were analysed using the CFX Manager software 3.1 (Bio-Rad, USA). Gene expression for each target gene was normalised to gene expression levels of chicken glyceraldehyde 3-phosphate dehydrogenase (GAPDH) and β-2-microglobulin (β2M). Relative normalised expression was calculated using the 2−ΔΔCt method and data are represented as fold change compared to control. Primer pairs used for the qPCR analysis are shown in **Table 1**.

**Table 1:**
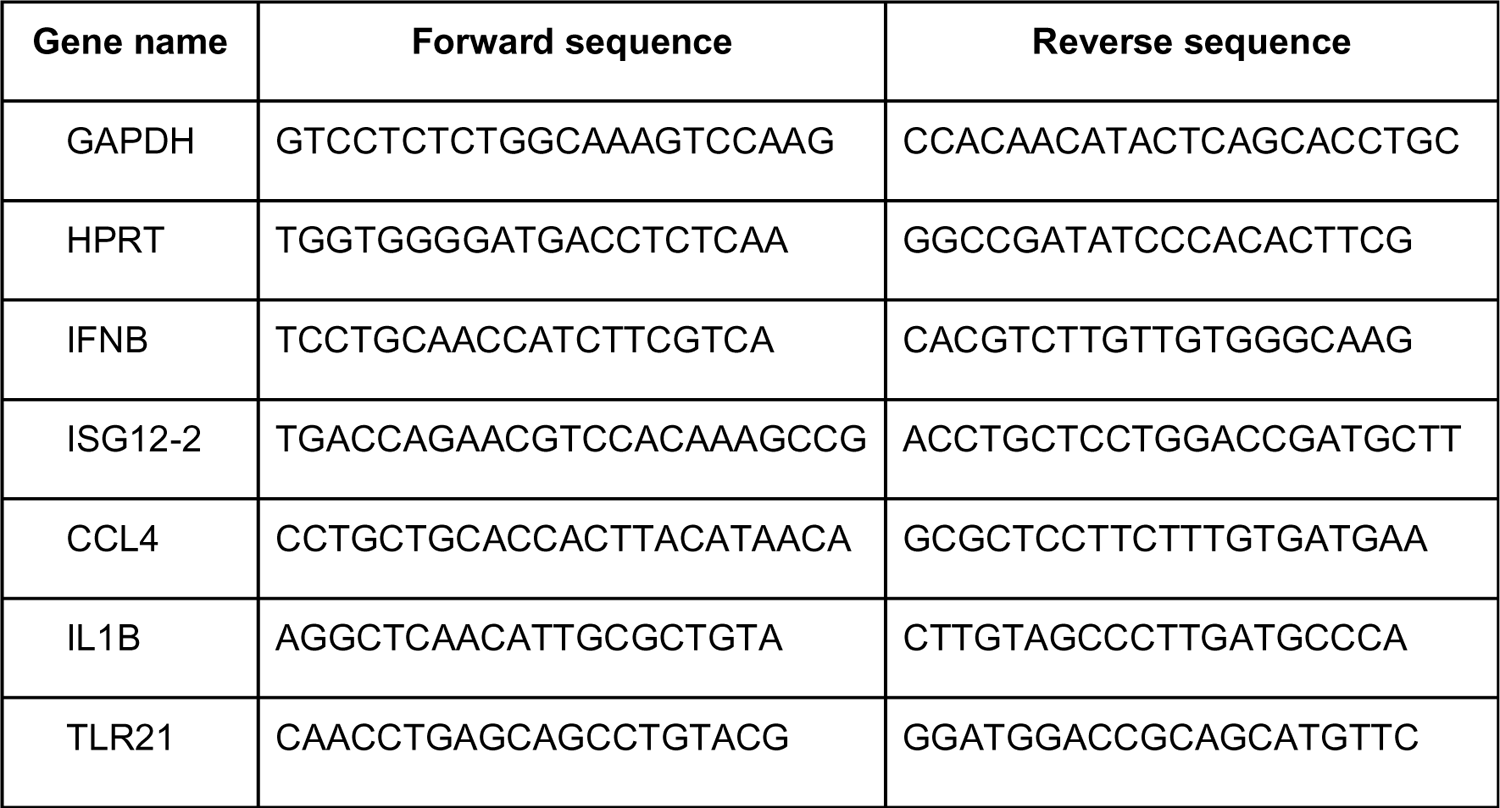
qPCR primer sequences.

### 2.7 Cell Viability

HD11 viability following different stimuli was assessed using the fluorescent DNA intercalator 7-aminoactinomycin D (7-AAD, BD Biosciences, USA). 7-AAD assay allows identifying populations representing live, apoptotic, and late apoptotic/dead cells. Briefly, following stimulation or transfection, supernatants were discarded, and the cells were harvested and washed in PBS. Cells were stained according to the manufacturer’s protocol and the viability was analysed by flow cytometry (BD FACS Calibur). Data were expressed as the percentage of 7AAD positive cells over total acquired events (50,000 cells).

### 2.8 Statistical Analysis

Prism 7 (GraphPad) was used to generate graphs and perform statistical analysis. Data were analysed using an unpaired t test with Welch’s correction unless stated otherwise. Data with P < 0.05 was considered significant and 2-tailed P-value were calculated and presented as: *p < 0.05, **p < 0.01, ***p < 0.001. Each experiment has at least two biological replicates unless stated.

## 3 Results

### 3.1 TLR21 is present across Aves and is closely related to TLR13

Several vertebrate TLRs have the ability to sense and respond to nucleic acids. TLRs 3, 7, 8, 9, 13, 19, 21 and 22 have all been reported to detect specific DNA or RNA species. Most nucleic acid-sensing TLRs bind to RNA, and only TLRs 9 and 21 have been shown to detect DNA, specifically DNA that contains CpG motifs. There is some recorded functional redundancy between TLR9 and TLR21 and some vertebrate genomes that contain both genes (Keestra et al., 2010; Temperley et al., 2008). In birds including chickens, the presence of TLR21 but not TLR9 is recorded, suggesting that TLR21 is the dominant CpG sensor. Some birds, however, have no annotated TLR21 or TLR9 in their genomes, but are noted to contain TLR13 genes. Since TLR13 is the closest related TLR to TLR21 (Temperley et al., 2008), we considered the possibility that some of bird TLR21 genes are incorrectly annotated as TLR13. To understand further the evolutionary relationships between the TLR13/21/22 family (which also contains TLR11/12 that sense profilin and flagellin) we conducted a maximum likelihood phylogenetic reconstruction of this family across vertebrates (Fig 1A). The bird TLR21s cluster in this tree in a group closest to reptile TLR21s and within a well-supported clade containing all vertebrate TLR21s, clearly separate from the other family members. This analysis clearly indicates that aves genomes only contain TLR21 from this family and do not encode TLR11/12/13/22. As such and considering the lack of any identifiable TLR9 genes in bird genomes, we followed the hypothesis that birds express only one CpG DNA sensing TLR, namely TLR21.

**Figure 1.**
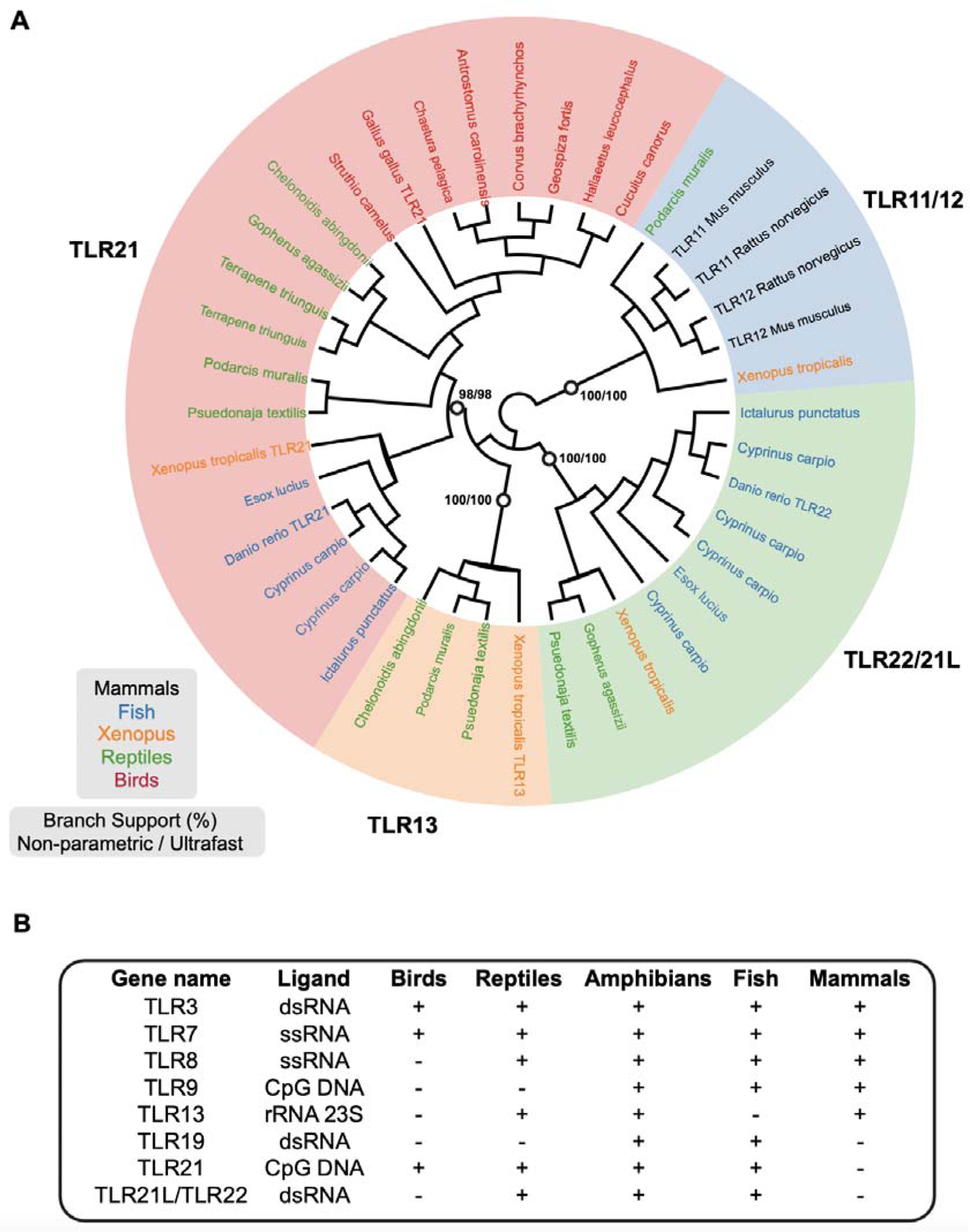
Phyletic distribution of nucleic acid sensing TLR families. (A) Maximum likelihood reconstruction of the closely related TLR11/12/13/21/22 families from vertebrates. Support for key branches is shown (open circles, non-parametric/ultrafast bootstrap support). Taxa are coloured by phylogeny (see key). The major clades separating TLR11/12, TLR21, TLR13, and TLR22/21L families are robustly supported. All avian sequences, annotated as either TLR21 or TLR13, group together within the TLR21 clade. (B) Distribution of nucleic acid sensing TLRs (‘-’ indicates not present, ‘+’ indicates present). Data assembled from published analyses and our own analysis in A.

Out of all vertebrates, birds express fewer nucleic acid sensing TLRs, encoding only TLR3 and TLR7 as RNA-sensing TLRs, and TLR21 as a DNA-binding TLR. This is in contrast to amphibians and fish whose genomes encode 7-8 nucleic acid sensing TLRs (Fig. 1B). It is possible, therefore, that bird TLRs have evolved to respond to a broader range of ligands than those of other vertebrates, that the function of other vertebrate TLRs is encoded in other innate immune mechanisms in birds. Since TLR21 is the only avian DNA binding TLR, its signalling mechanisms and outputs have broad consequences for avian innate immunity, immune regulation and vaccinology.

### 3.2 CpG DNA triggers a type-I interferon (IFN-I) response in chicken macrophages

We assessed the ability of chicken macrophages to respond to CpG DNA using a Class B CpG ODN containing a full phosphorothioate backbone. ODN2006 is known to activate mammalian TLR9 and avian TLR21 and to trigger an inflammatory transcriptional programme (Chuang et al., 2020; Keestra et al., 2010). We carried out a dose response stimulation of ODN 2006 in a chicken macrophage line and observed by qPCR dose-dependent transcription of type-I interferon (*IFNB*) and the chemokine *CCL4* (Fig. 2A). In mammals *IFNB* transcription is dependent on co-activation by the transcription factors IRF3 and NF-κB acting in synergy on the same *IFNB* promoter in a complex known as the enhanceosome (Thanos and Maniatis, 1995). CCL4 on the other hand, is mainly an NF-κB responsive gene in mammals (Rezzonico et al., 2001). These data indicate that TLR21 can potently activate a transcriptional IFN-I and chemokine response in chicken macrophages. Next, we tested the response of HD11 cells to several nucleic acid species in parallel and assessed the transcription of four genes, *IFNB*, *CCL4*, *IL1B* and the interferon-stimulated gene *ISG12.2* (encoding IFI6). ODN induced all four genes to a higher level of expression than intracellular DNA stimulation or poly(I:C) transfection, apart from *ISG12.2* that was induced to the greatest level by intracellular DNA, sensed by the cGAS/STING pathway (Fig. 2B). Indeed, we have previously shown that, consistent with mammalian cGAS mechanism, stimulation of HD11 cells with DNA species or 2’3’-cGAMP resulted in robust IFN-I transcription, dependent on the cGAS/STING pathway (Oliveira et al., 2020). We did not observe IFNLJ, IFNλ or IFNβ transcription by HD11 cells in response to ODN indicating the primary interferon response to CpG DNA in chickens is production of interferon beta.

**Figure 2.**
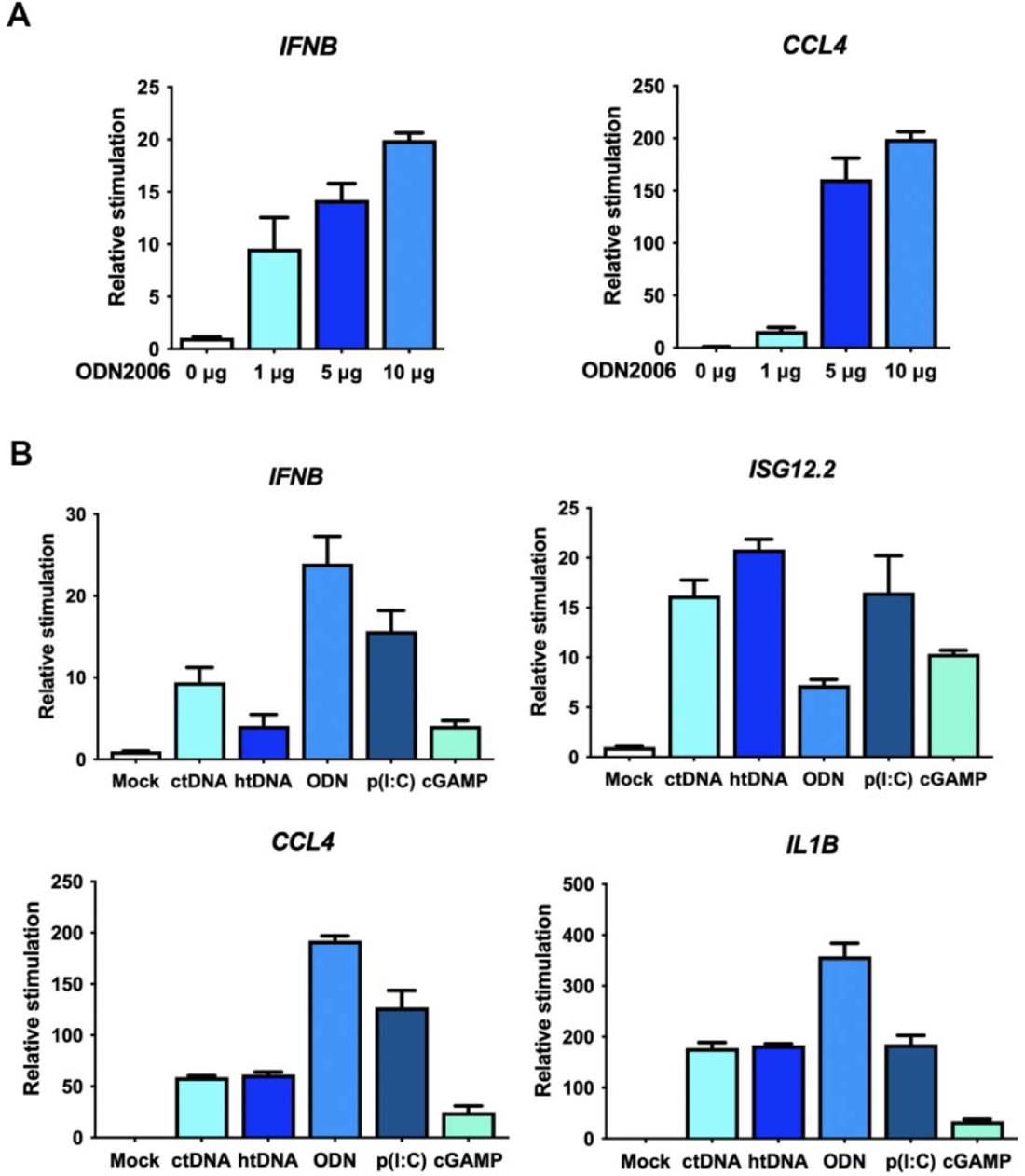
CpG DNA drives a type-I interferon response in chicken macrophages. (A) HD11 cells were stimulated with the indicated doses of type B CpG DNA (ODN2006) and transcription of *IFNB* and *CCL4* was measured by qRT-PCR 6 h later. (B) HD11 cells were stimulated with the indicated PRR ligands (incubation with ODN2006 or Poly(I:C), or transfection with htDNA, ctDNA or 2’3’-cGAMP) and transcription of the indicated genes was measured by qRT-PCR 6 h later.

Previously we have shown that pre-stimulation with IFNLJ can boost the transcriptional response to nucleic acid stimulation (Oliveira et al., 2020). ODN-driven *IFNB* transcription was also enhanced by pre-stimulation with interferon alpha (Fig. 3A) and the transcription of *TLR21* itself was upregulated by a combination of ODN2006 and IFNLJ but not by individual stimuli (Fig. 3B). These data indicate that there is a positive feedback loop downstream of CpG DNA that activates *IFNB* transcription and can enhance TLR21 expression and signalling in chicken macrophages.

**Figure 3.**
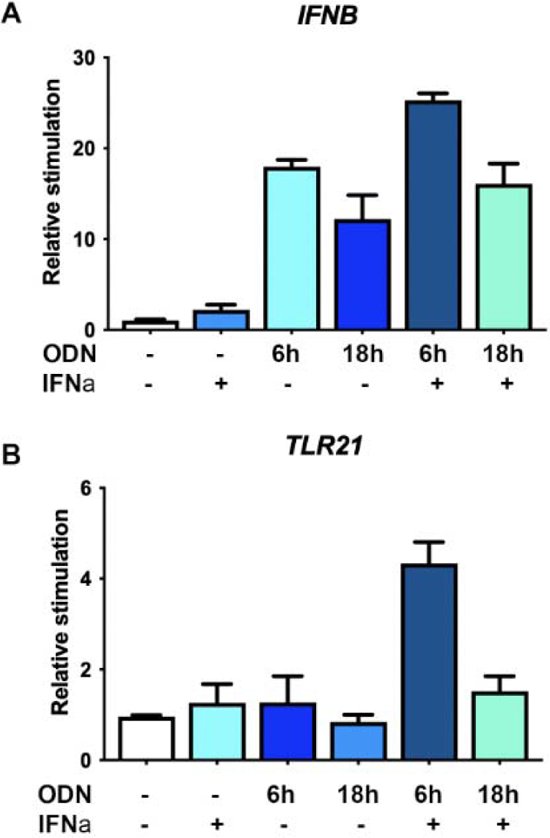
TLR21 expression is enhanced by the synergistic activity of IFN-I and CpG. HD11 cells were preincubated with chicken interferon alpha (50 ng/ml for 16 hours) before being stimulated with type B CpG DNA (ODN2006) for the indicated times. Transcription of *IFNB* and *CCL4* was subsequently measured by qRT-PCR.

### 3.3 Interferon production by TLR21 is partially NF-kB dependent

Next, we analysed the dependence of nucleic acid-dependent transcriptional outputs on NF-κB activity. A HD11 reporter cell was used to show that, when stimulated with a range of PAMPs, those stimuli that specifically activate TLR signalling are potent activators of NF-κB activity (Fig. 4A) and this signal can be inhibited with the small molecule compound BAY-11-7082 that blocks IKKLJ/β activity, but not with BX795 that inhibits TBK1/IKKε activity (Fig. 4 A,B). As such CpG DNA and LPS, but not non-CpG DNAs or RNAs, can efficiently signal via NF-κB.

**Figure 4.**
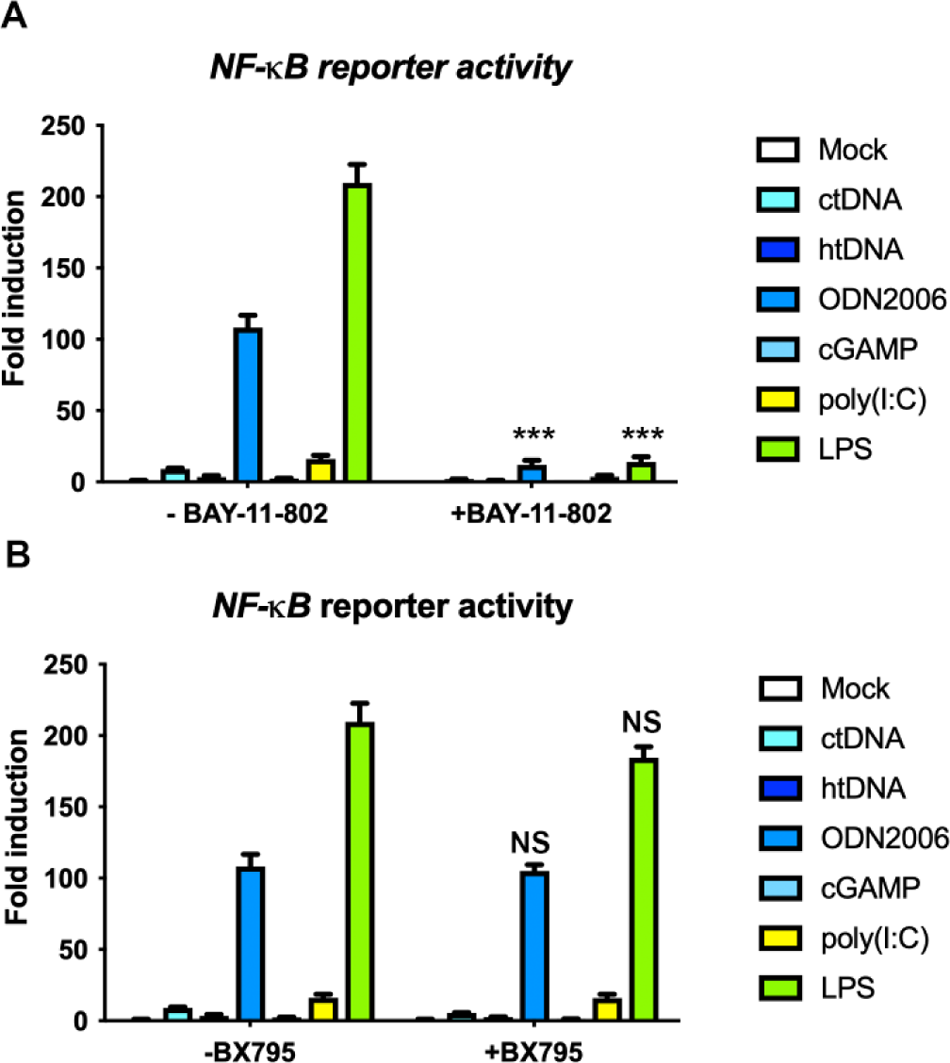
CpG DNA triggers potent NF-kB activation. HD11 NF-κB reporter cells were stimulated with the indicated PAMPs in the presence and absence of (A) BAY-11-7082 or (B) BX795, and 24 hours later luciferase activity was measured. NS-non significant, **p < 0.01, ***p < 0.001, n≥3.

Using both BAY-11-7082 and BX795 we dissected the relative contribution of NF-κB versus TBK1-dependent signalling on the transcription of *IFNB*, *ISG12.2, CCL4* and *IL1B* downstream of CpG and non-CpG containing DNAs (Fig. 5). We observed that *CCL4* and *IL1B* transcription was abrogated using the IKKLJ/β inhibitor, indicating that these two genes are almost exclusively transcribed in an NF-κB dependent manner downstream of DNA stimulation (with or without CpG motifs) In chickens. *IFNB* transcription was partially inhibited by BAY-11-7082 following ODN stimulation, consistent with its transcription being partially NF-κB dependent. *ISG12.2* transcription was inhibited neither by BAY-11-7082 nor BK795 in response to CpG DNA, indicating that ODN-driven IFN-I responses in chickens are not dependent on TBK1/IKKε or NF-κB (Fig. 5). These data show that there are two distinct signalling arms downstream of CpG DNA stimulation. One signalling arm drives inflammatory gene expression via NF-κB and is responsible for *IL1B* and *CCL4* transcription and one arm that drives a TBK1/IKKε-independent pathway responsible for *ISG12.2* transcription. These two pathways combine to drive IFNB transcription in response to CpG DNA.

**Figure 5.**
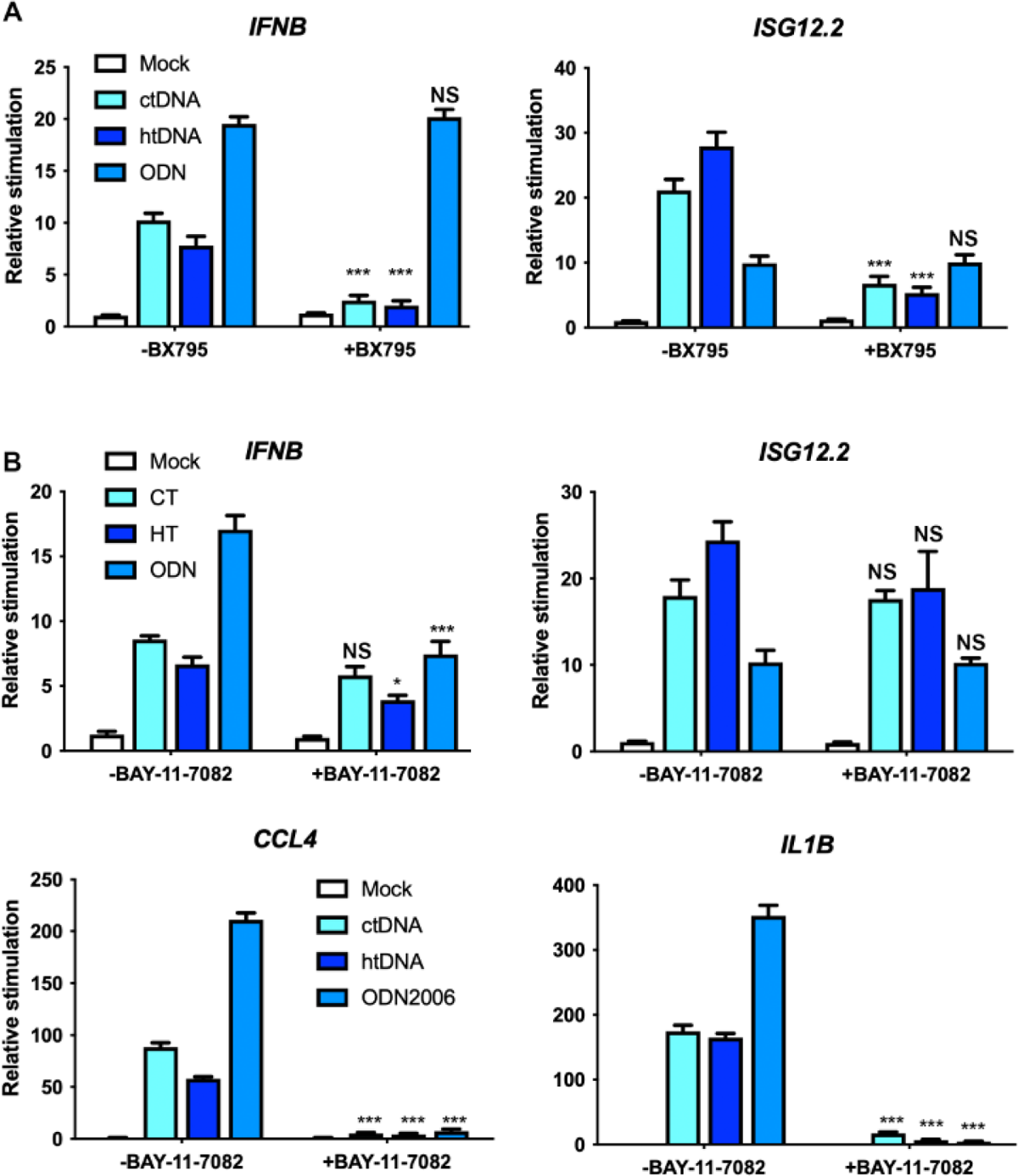
CpG-driven interferon production is independent of TBK1. HD11 cells were stimulated with ODN2006, or by transfection with htDNA or ctDNA, in the presence or absence of (A) BX795 or (B) BAY-11-7082. Transcription of the indicated genes was measured by qRT-PCR 6 h later. NS-non significant, **p < 0.01, ***p < 0.001, n≥3.

### 3.4 TLR21 and IRF7 are required for CpG sensing in chickens

To determine the contribution of TLR21 to CpG sensing we used CRISPR/Cas9 to generate clonal TLR21 knockout HD11 cells (Fig. 6A). Stimulation of these cells demonstrated that TLR21 is absolutely required for *IFNB*, *ISG12.2*, *CCL4* and *IL1B* transcription in response to CpG DNA (ODN2006) stimulation but not in response to cytoplasmic (non-CpG) DNA stimulation. This confirms that TLR21 is the CpG DNA sensor in chickens and drives inflammatory and IFN-I responses.

**Figure 6.**
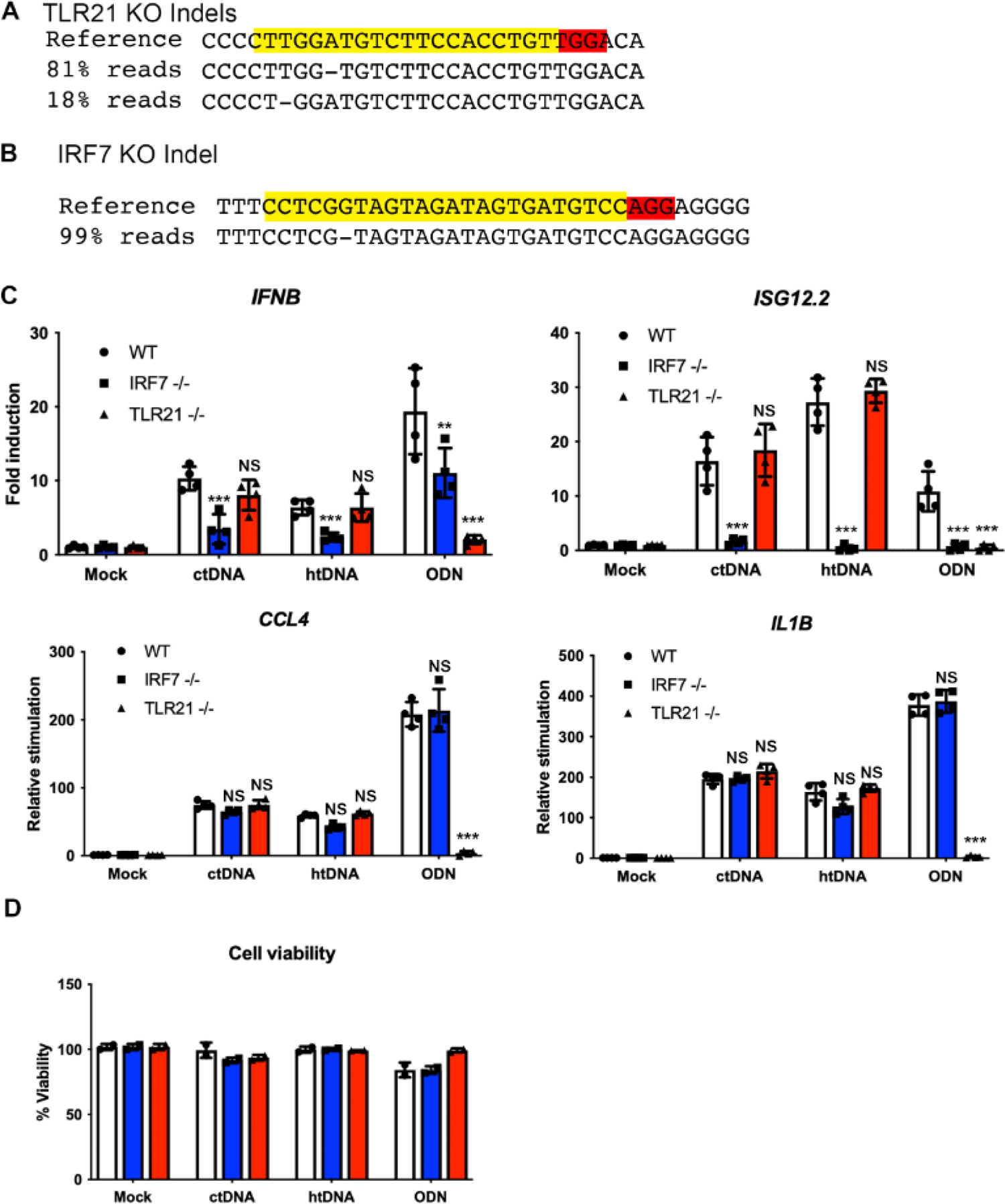
TLR21 is the primary CpG DNA sensor and signals via IRF7 in chickens. (A) TLR21 knockout and (B) IRF7 knockout HD11 cells were generated by CRISPR/Cas9 editing and sequenced to define indels targeted by the gRNAs (yellow) including PAM sequences (red). (C) WT, TLR21 KO and IRF7 KO HD11 cells were stimulated with ODN2006, or by transfection with htDNA or ctDNA and transcription of the indicated genes was measured by qR T-PCR 6 h later. (D) WT, TLR21 KO and IRF7 KO HD11 cells were stimulated with ODN2006, or by transfection with htDNA or ctDNA and cell viability measured 24 hours later. NS-non significant, **p < 0.01, ***p < 0.001, n≥3.

Since ISG12.2 transcription downstream of CpG DNA was independent of NF-kB activity (Fig. 5), we postulated that activation of this gene was dependent on a pathway that uses TLR21 and IRF7. IRF7 in chickens has been proposed to act in a similar manner to mammalian IRF3, as an activator of IFN-I downstream of key antiviral signalling pathways (Cheng et al., 2019; Santhakumar et al., 2017), and TLR9 in mammals can activate IFN-I responses in dendritic cells specifically via a pathway that requires IRF7 (Honda et al., 2005b). We found that the presence of TLR21 is phylogenetically correlated with IRF7 in avian genomes (Supplementary Figure 1). Further, analysis of the amino acid sequences at the C-terminus of avian IRF7s indicated the presence of serine residues that are conserved in both mammalian IRF3 and IRF7 including GASSL/LSSA and SNSHPLSLTS motifs (Supplementary Figure 2) (Caillaud et al., 2005; Chen et al., 2008). This conservation of functional residues suggested that chicken IRF7 may be able to carry out the functions of both mammalian IRF3 and IRF7, and possibly work downstream of TLR21 to drive IFN-I responses to CpG DNA. To assess this, we generated IRF7 knockout HD11 cells (Fig. 6B). Stimulation of these cells with intracellular DNA and ODN did not impact the transcription of *CCL4* or *IL1B,* but abrogated *ISG12.2* transcription and reduced *IFNB* transcription (Fig. 6C). These data define that IFNB and ISG12 are canonical markers of nucleic acid detection in avian macrophages upon binding of the latter to the two principal PRR pathways sensing dna, namely TLR21 and cGAS/STING. Importantly none of these stimuli caused significant changes in cell viability (Fig. 6D). Overall these data are consistent with a model where TLR21 drives both NF-kB and IRF7 activation and these transcription factors combine to generate a IFN-I and proinflammatory response to CpG DNA in chickens (Fig. 7).

**Figure 7.**
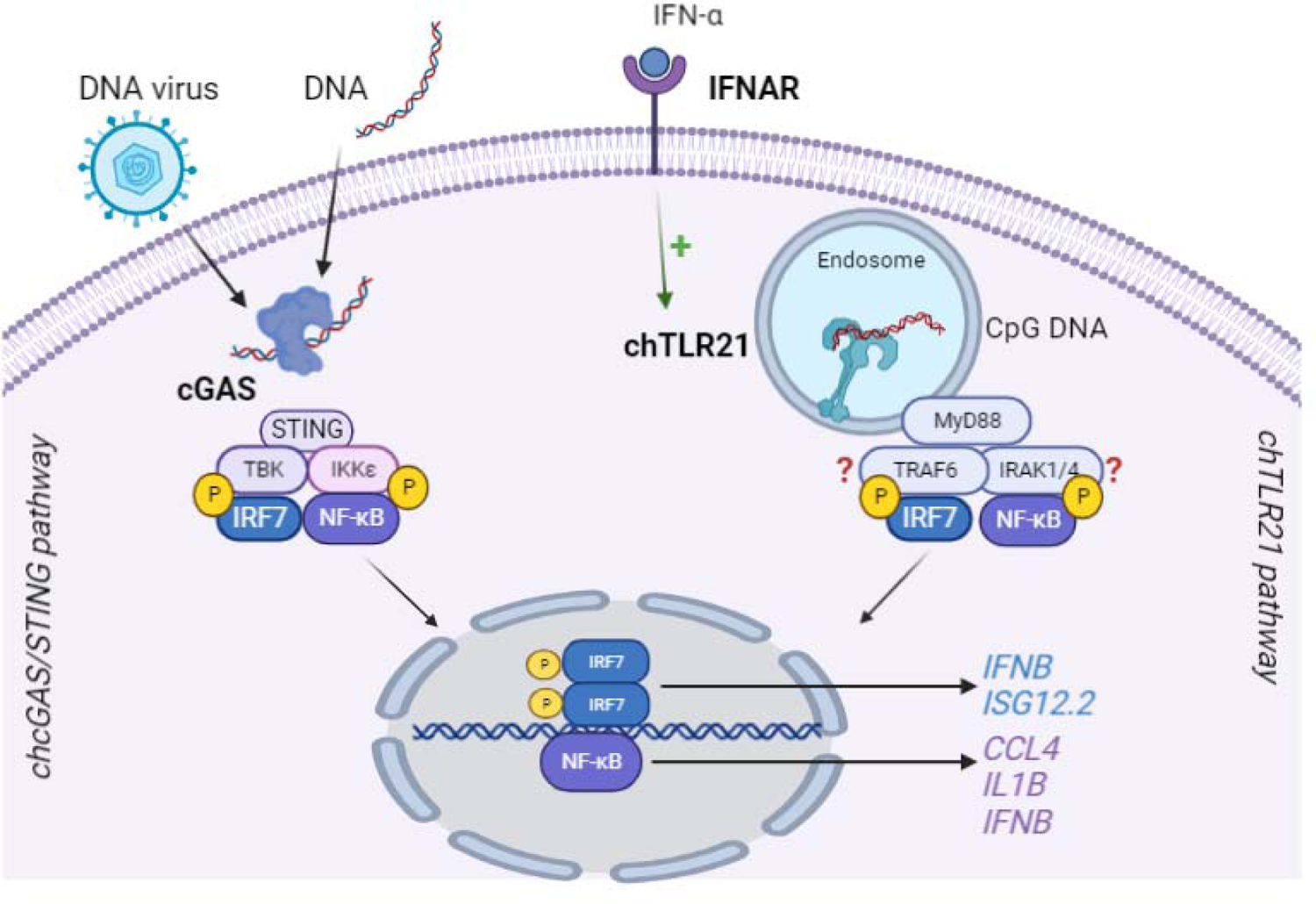
TLR21 and cGAS sensing of DNA in chicken macrophages. Schematic diagram outlining sensing and downstream signalling of avian DNA sensing PRRs. Sensing by TLR21 or cGAS results in activation of IRF7 and NF-kB that drive specific or synergetic transcription of inflammatory and antiviral target genes. IRF7, for example, activates ISG12.2, NF-kB activates *CCL4* and *IL1B* and together IRF7 and NF-kB drive *IFNB* transcription.

## 4 Discussion

Avian TLR21 has evolved to sense and respond to CpG DNA and is hence an important anti-microbial PRR. Here we confirm that the TLR21 is the main CpG DNA sensor in chickens. The ability to detect CpG DNA was lost in cells where TLR21 had been knocked out. This is in keeping with previous papers using siRNA approaches to functionally analyse the contribution of TLR21 to innate sensing of DNA (Brownlie et al., 2009; Keestra et al., 2010). We found that TLR21 can robustly activate an IFN-I response following stimulation with CpG DNA. This response is dependent on downstream signalling via both NF-κB and IRF7, key transcription factors in the innate immune response. NF-κB and IRF7 act synergistically to induce *IFNB* transcription and independently to induce expression of proinflammatory (*CCL4* and *IL1B*) and antiviral genes (*ISG12.2*). This response differs from the sensing of intracellular DNAs as it is independent of TBK1/IKKε, the kinases normally associated with IRF activation downstream of nucleic acid sensing PRRs (Balka et al., 2020). Instead, TLR21 signalling is more similar to the mammalian TLR9 mechanism of activating IFNa transcription in dendritic cells. Here, TLR9 activates both NF-κB and IRF7 via a signalling complex that includes IRAK4 and TRAF6 (Bryant et al., 2015; Honda et al., 2005a, 2004). As such chTLR21 signals in a similar manner to TLR9, despite the evolutionary distance between TLR21 and TLR9. Liu *et al*. conducted phylogenetic analyses to investigate the orthologous and paralogous relationships among vertebrate Toll-like receptors (TLRs) and to identify TLR subfamilies (Liu et al., 2020). This analysis revealed that TLR21 is the sole member of the TLR13 subfamily found in avian species. TLR13 functions as a sensor for bacterial 23S ribosomal RNA. The absence of TLR13 in several mammalian species, including humans, has been attributed to various explanations. These range from the avoidance of autoimmune responses triggered by mRNAs displaying the same sequence as the immunostimulatory sequence in bacterial rRNA, to compensatory mechanisms involving other TLRs (Li and Chen, 2012). However, the immunological implications of losing both TLR9 and TLR13 genes in the ancestor of modern birds necessitate further investigation. Nonetheless, the profile of TLR21 signalling in birds strongly suggests a potential compensatory mechanism for the loss of TLR9.

Unlike mammalian IRF3, which is constitutively expressed and stable in most cells, mammalian IRF7 remains at low levels in most cell types, and has a very short half-life (Sato et al., 2000). Precise control over the expression and functioning of IRF7 holds utmost significance in orchestrating the optimal production of type-I IFN necessary for normal physiological processes reliant on IFN signalling. As chickens have lost IRF3, it is likely that chIRF7 replaces IRF3’s functions in PRR signalling in this species (Angeletti et al., 2020; Huang et al., 2010). Indeed our data here, showing that chIRF7 is required for htDNA- and ctDNA-driven *IFNB* and *ISG12.2* transcription is consistent with previous studies showing that chIRF7 functions in the STING and MAVS intracellular nucleic acid sensing pathways (Cheng et al., 2019). We also find here that chIRF7 is also required for TLR21 signalling analogous to mammalian IRF7 in the TLR9 signalling pathway. As such it appears that chIRF7 has evolved to carry out at least some of the combined functions of mammalian IRF3 and IRF7. The fact that chickens (and galliforms in general) do not express an IRF3 highlights the relevance of investigating regulatory mechanisms upstream its activation upon pathogen challenges. Despite the remarkable advancements made in the last decade numerous unresolved questions persist concerning IRF7 regulation of type-I IFN responses in birds (Kim et al., 2020; Wang et al., 2021).

CpG-B ODNs, classified as Class B (type K), are linear oligodeoxynucleotides ranging from 18 to 28 nucleotides in length. They possess a phosphorothioate backbone and incorporate one or more 6-nucleotide CpG motifs. Previous studies showed that mammalian TLR9s have preferences in terms of recognizing different CpG-ODN nucleotide sequences (Yamamoto et al., 2000). The ideal motif for human CpG-B ODNs is GTCGTT, while for mice, it is GACGTT (Chuang et al., 2020). However, chicken and duck TLR21s, unlike mammalian TLR9s and TLR21 from other taxa, do not distinguish between different types of CpG-hexamer motifs, indicating that CpG-ODNs ranging from 15 to 31 nucleotides can activate avian TLR21s regardless of the spacing between the two GTCGTT. While the CpG-ODNs with the AACGTT motif are not well studied in birds, those with the CTCGTT or GACGTT motifs have already shown adjuvant activity (Lee et al., 2010). The broad CpG-ODN sequence recognition profile exhibited by chicken and duck TLR21 implies a wide range of CpG-ODN options that can be used as adjuvants in vaccines or as immune-modulators (Bavananthasivam et al., 2018; De Silva Senapathi et al., 2018; Gaghan et al., 2023; Scheiermann and Klinman, 2014). Moreover, CpG-ODN can be administered to chickens through different routes, including oral, intranasal, subcutaneous, and *in ovo* injections (De Silva Senapathi et al., 2018; Talebi and Arky-Rezai, 2019). This discovery enhances the potential for selecting optimal CpG-ODNs to effectively enhance antigen-dependent immune responses in poultry birds (Gupta et al., 2014). As immunomodulatory approaches become increasingly integral to large-scale poultry production, it becomes crucial for future research to delve into the underlying mechanisms and modes of action that drive these outcomes.

## Concluding remarks

Studying the relationships between avian TLR21 and IFN-I production outputs is crucial to elucidate CpG-ODN-driven immune responses in poultry species. To our knowledge, this is the first work to depict the relative contribution of the main transcription factors involved in chTLR21 transcriptional outputs, namely IRF7 and NF-κB. Since avian TLR21s exhibit broad recognition of various CpG-ODN sequences, the innate immune system of poultry birds possesses a robust ability to detect DNA-associated molecular patterns from microbes. Therefore, the wide range of CpG-ODN options available for use as immune-stimulatory agents and a better definition of immunological outputs downstream avian TLR21 activation will widen its application as vaccine adjuvants compared to other mammalian livestock species.

## Acknowledgements

This work was funded by BBSRC grant BB/S001336/1 (BJF and CEB) and by EUROFERI FEDER-FSE grant N° EX 010233 (RG and ST).

---

**Supplementary Figure 1.**
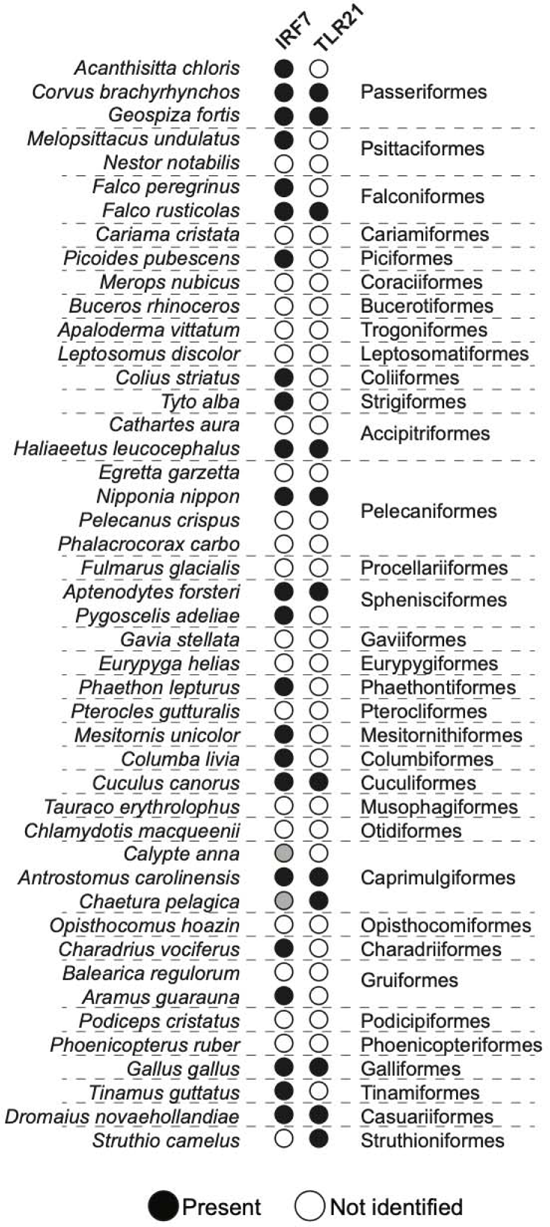
Phyletic distribution of TLR21 and IRF7 across birds. Overview of presence or absence of TLR21 and IRF7 genes in birds. Annotation of TLR21 sequences was confirmed by phylogenetic analysis, annotation of IRF7 sequences was confirmed by reciprocal BLASTp against human sequences. Open circles denote no corresponding sequence identified and inferred absence, closed circles denote presence.

**Supplementary Figure 2.**
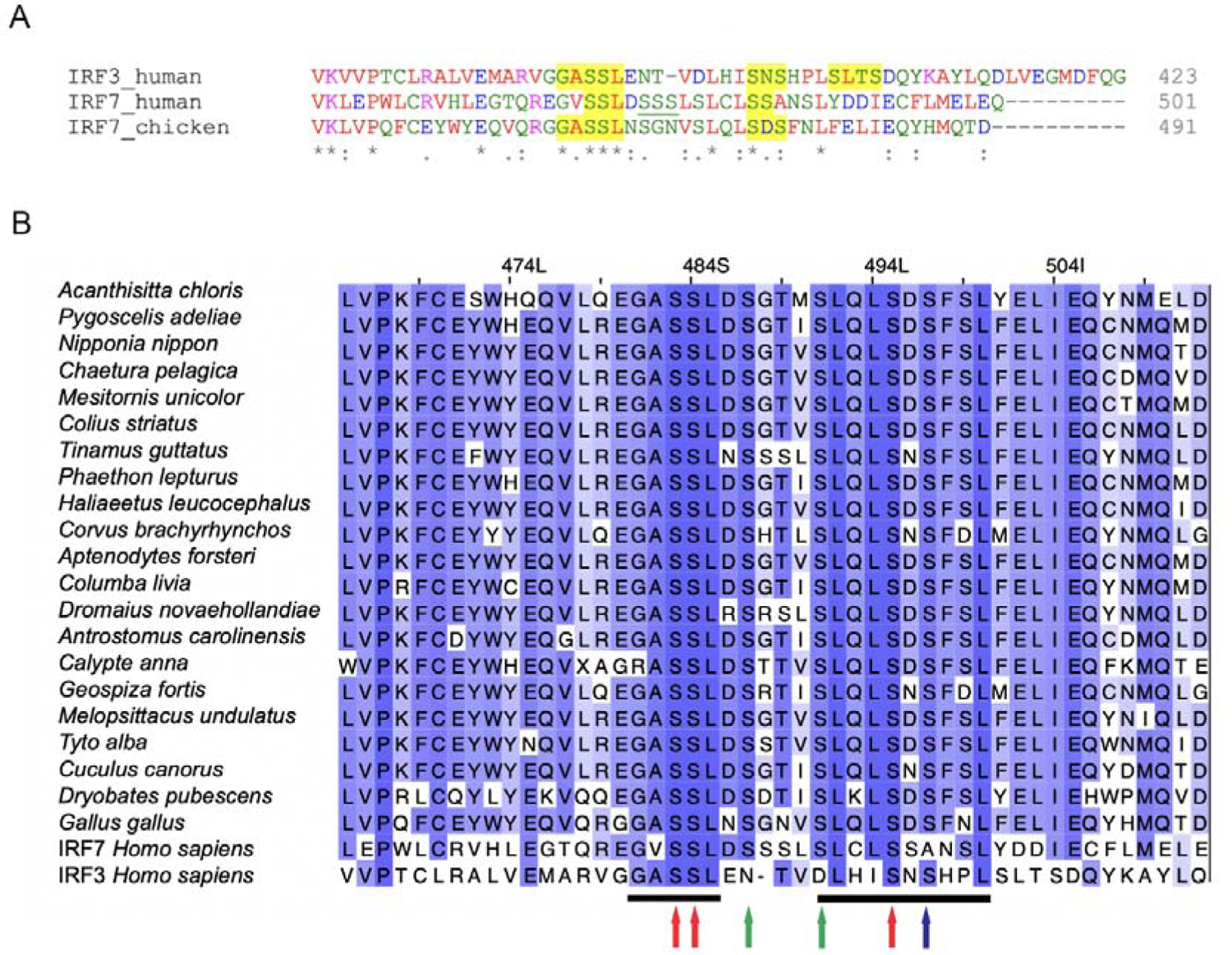
Mammalian IRF3 and IRF7 phosphorylation sites are present in avian IRF7s. A) Amino acid alignments of the C terminal regions of human IRF3 and IRF7 with chicken IRF7. Highlighted in yellow are the key phosphorylation sites in mammalian IRFs that contain multiple serine residues essential for IRF activation and dimerisation. B) Alignment of multiple avian IRF7s with human IRF3 and IRF7 indicating complete conservation of key serine residues in bird IRF7 that are essential for human IRF3 activation (blue arrows), human IRF7 activation (green arrows) or both (red arrows).

**Supplementary Table 1:**
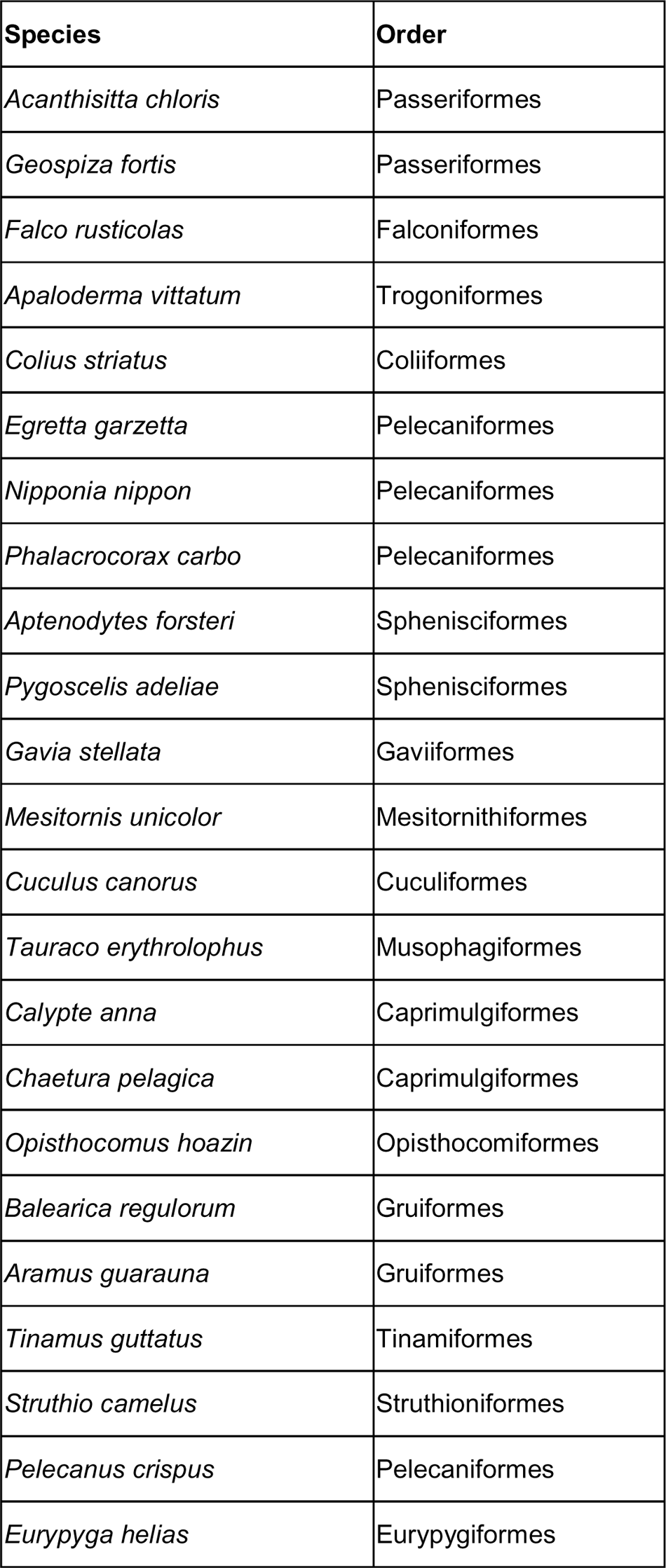
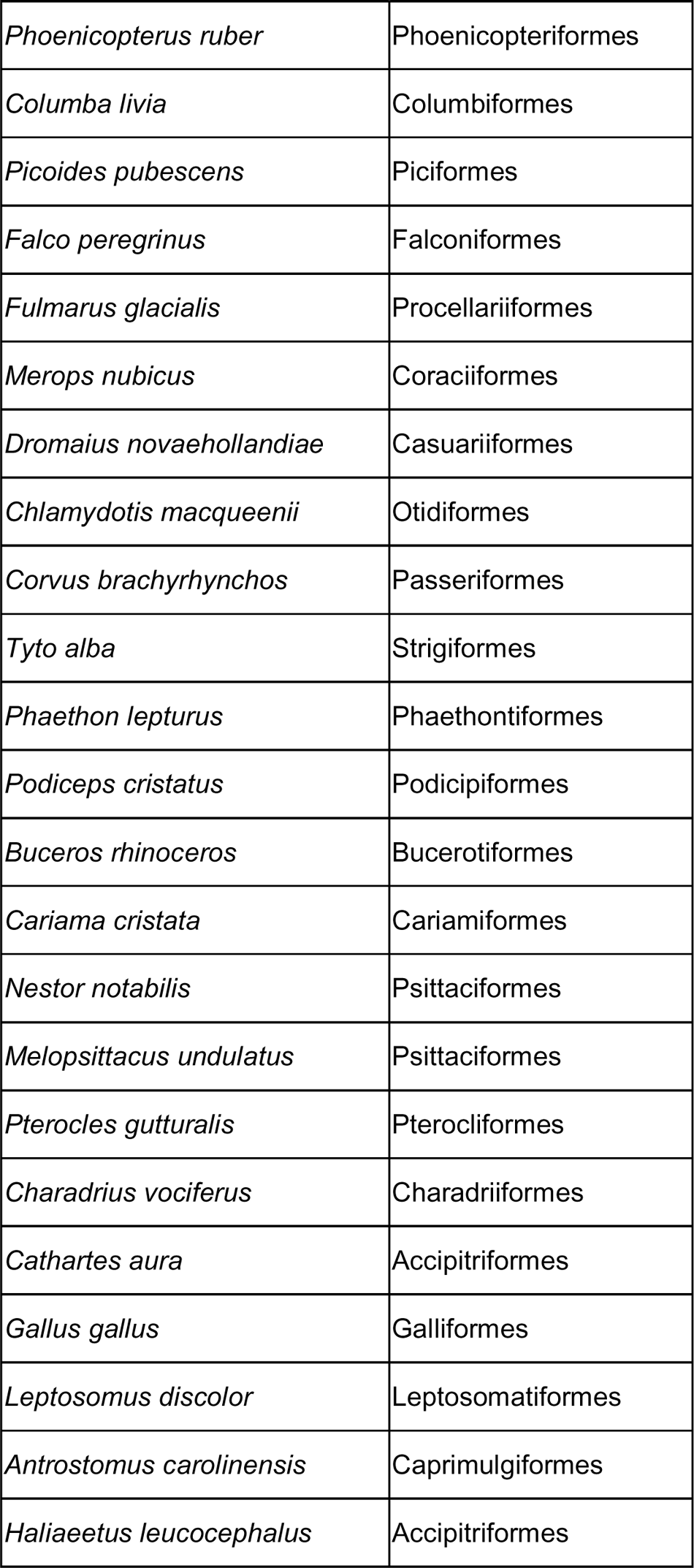
Bird species analysed in this study.

